# Systematic molecular glue drug discovery with a high-throughput effector protein remodeling platform

**DOI:** 10.1101/2025.07.17.664724

**Authors:** Jia Lu, Kate Stuart, Rebecca Teague, Chloe Tarry, Robert Yan, Tabitha Morgan, Elizabeth Nock, Darcie S Mulhearn, Matt Jones, Michael H Knaggs, Ruben Alvarez Fernandez, Willie Yen, Aris Aristodemou, Eleanor Thompson, Penny Hayward, Juliane F Ripka, Bethany C Atkinson, Abigail Dear, Aruba Farooq, Mat Calder, Miguel B Coelho, Laura R Butler, Alberto Moreno, Christian Dillon, Richard J Boyce, Benedict CS Cross

## Abstract

Realising the promise of new medicines that operate through a targeted molecular glue-induced degradation mechanism requires systematic tools that can uncover the relevant principles of neomorphic protein-protein interactions. Whilst some monovalent glue degraders have been found through serendipity, the rules for small molecule attributes and the pairs or complexes of proteins that are amenable to drug-induced proximity control remain poorly articulated. Here we introduce a new approach to address this by using programmed libraries of intramolecularly edited proteins to expand protein surface landscapes and trigger new druggable interactions. We show that effector proteins, such as the E3 ligase Cereblon, can be engineered to provoke neomorphic activity by inducing the degradation of new client proteins and that these *de novo* interactions provide a blueprint from which new small molecule degraders can be built. As a demonstration of the approach, we use the platform to identify new non-IMiD molecular glue degraders of the oncology target GSPT1.

**SUMMARY:**

- Molecular glues are a highly important and promising new form of therapeutic agent, but rationalising their discovery has so far been impossible
- GlueSEEKER screening enables prospective monovalent drug discovery by using high-throughput deep mutational scanning to re-engineer the function of effector proteins like E3 ligases
- We used this approach to enable the computational discovery of small molecule glues which degrade the oncology target GSPT1 and show how the technology can be used across new targets

## INTRODUCTION

Protein-protein interactions (PPIs) provide an essential mechanism to control regulatory circuits in biology and are a crucial node where disruption can cause pathogenicity in diseases from cancers to neurodegenerative decline. Perturbation of toxic PPIs can be a valuable route to therapy, and whilst some progress towards targeted medicine development has been made (Lu et al., 2020), the canonical occupancy-driven mode-of-action (MoA) renders some important therapeutic targets ‘undruggable’ or without suitable pockets for binding high-affinity ligands. Induction of PPIs is emerging as a novel strategy, including the use of targeted protein degradation (TPD) as a gain-of-function drug modality. In contrast to inhibitors, TPD causes the removal of pathogenic targets using the cell’s own waste disposal system. Because of the event-driven MoA, TPD has the potential to overcome limitations of small molecule inhibitors, such as drug resistance, selectivity and poor tolerance (Tsai et al., 2024). PROteolysis-TArgeting Chimeras (PROTACs) and molecular glue degraders (MGDs) are two major modalities of TPD, both of which drive interaction between an E3 ligase and a protein of interest (POI). MGDs often possess more favourable pharmacokinetic properties than PROTACs and are a focus of substantial interest across drug discovery efforts, yet their development has been hampered by a lack of tools which can help determine the common chemical features and design properties that define them. MGDs can have minimal measurable binary affinity for the participating individual proteins making the determination of suitable screening assays challenging. Promiscuity of activity is also a challenge, where MGD activity risks potential off-target effects or masks on the on-target activity. Consequently, most MGDs are discovered only serendipitously and most often are based on analogues of well-established ligands that bind the E3 ligase Cereblon (CRBN) as the predominant effector protein.

In order to explore the vast promise of MGDs, we sought to revisit the fundamental properties of the modality using a systematic screening approach. We posit that since MGDs are defined by their ability to mimic the features of PPI, it follows that neo-PPI will report on the chemical requirements for an MGD. We constructed a synthetic, cell-based screening platform that uses insertion mutagenesis as a deep mutational scanning application to edit the surfaces of effector proteins such as E3 ligases. By surveying the resulting degradation landscape of these edits we can identify the acquisition of neosubstrates using high-throughput pooled genetic screening and rationalize the chemical space required for inducing novel E3:POI interactions. The resulting maps are idealised pharmacophores for the design of small molecules that structurally and functionally mimic the neomorphic E3:POI interaction. We call this new approach GlueSEEKER and show its application through the discovery of new variants of CRBN which are able to degrade the oncology target GSPT1, a prototypical molecular glue target, and the translation of these unique observations to the discovery of cellularly active small molecule MGDs.

## RESULTS

### Neomorphic gain-of-function of E3 ligases by protein editing on functional hotspots

To identify gain-of-function modifications to a nominated E3 ligase which can lead to induced degradation of neosubstrates, we first interrogated the substrate binding interface of the CRBN protein through insertional mutation-based perturbation. A series of short peptidic sequences were introduced in-frame within the coding sequence of the E3 ligase to determine suitability for large-scale screening. Epitope tags were introduced into two distinct sites on CRBN (P1 and P2) followed by ectopic expression in wild-type CRBN-deleted cells which achieved comparable protein expression to endogenous protein (not shown) and was used to examine the functional conservation and any neomorphic activity of the ligase.

To test if the modified CRBN could exert typical ligase activity even when short non-native sequences were introduced into the protein sequence, we treated reconstituted cells with a heterobifunctional BRD4 degrader dBET6 (Winter et al., 2017). dBET6 induces the degradation of BRD4 in HAP1 parental cells (**Fig 1a**, lane 6) but not in HAP1^*CRBN-*^cells (lane 7). Reconstitution of CRBN activity by exogenous expression of CRBN_WT restores BRD4 degradation (lane 8). dBET6 was still able to induce BRD4 degradation of a CRBN_HA mutant at the P1 site (lane 9), albeit with modestly reduced effect compared to WT protein, and importantly degradation is inhibited by treatment with the neddylation inhibitor MLN4924 (lane 14). However, despite an HA insertion at the P2 site showing similar CRBN expression to P1, there is no observable BRD4 degradation (lane 10 and not shown), indicating that either enzyme function or the binding of the PROTAC molecule is compromised by the modification at this site in contrast to the P1 site.

**Figure 1.**
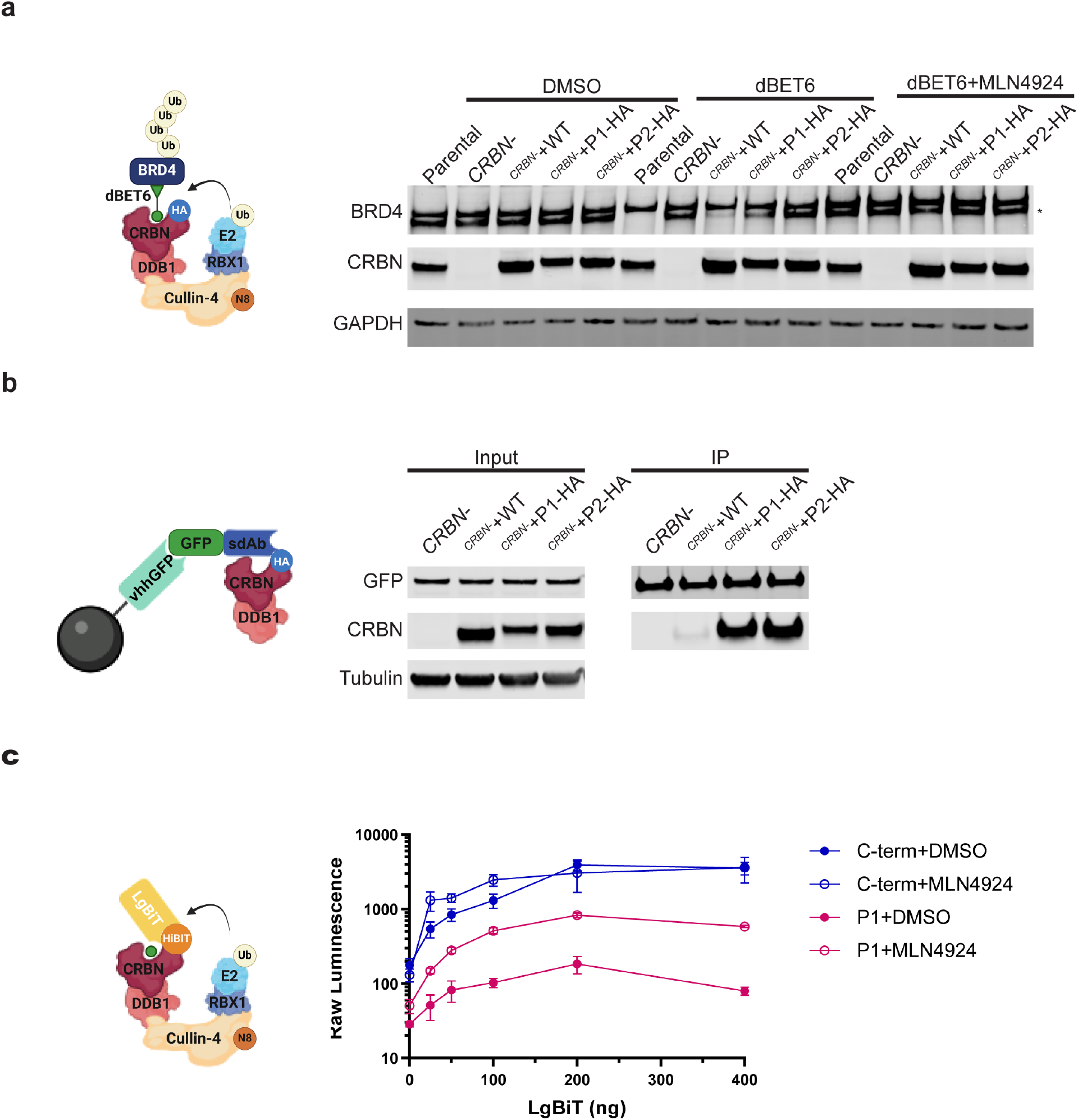
Modulation of E3 ligase activity via editing on functional hotspots. a) A surface-edited CRBN mutant is functional in inducing neosubstrate degradation similar to reported MoA. CRBN-overexpressing cells from b), together with HAP1 parental and untransduced HAP1^*CRBN-*^cells were treated with DMSO, dBET6 or dBET6+MLN4924 for four or 16 hours. Total cell lysates were harvested and analysed by western blot. dBET6-mediated degradation of BRD4 was observed in HAP1 parental, CRBN_WT and CRBN-P1 and is abolished by MLN4924 treatment. b) Surface editing of CRBN confers novel protein-protein interaction. An HA tag was inserted into two sites on surface of CRBN (P1 and P2), and HAP1^*CRBN-*^cells were transduced with lentivirus overexpressing CRBN_WT, CRBN-P1-HA or CRBN-P2-HA together with a GFP reporter fused to an anti-HA nanobody (HA-15F11) (Zhao et al., 2019). Using anti-GFP magnetic agarose, GFP-HA nanobody was enriched in the IP fraction. Anti-CRBN signal was only detected where HA tag was inserted in CRBN coding sequence. c) CRBN-P1-HiBiT is functional to induce neosusbtrate degradation. HAP1^*CRBN-*^cells were transduced with lentivirus encoding CRBN with a HiBiT tag inserted at the P1 site. Stable cells were then transfected with plasmids expressing LgBiT, followed by DMSO or MLN4924 treatment for 24 hours post transfection. Cells were lysed and luciferase activity was measured. Compared to CRBN containing a C-terminal fusion of HiBiT tag, a LgBiT dose-dependent reduction of luminescence signal was observed and was partially restored by MLN4924 treatment.

The ability of the CRBN_HA variant to induce a neomorphic PPI was examined by co-expression of the variants with an HA-nanobody reporter fused to a GFP molecule (Zhao et al., 2019). Co-immunoprecipitation from whole cell lysates using anti-GFP magnetic agarose showed that CRBN_HA was enriched in the co-IP fraction whilst the CRBN_WT was not (**Fig 1b**), indicating that the inserted HA tag confers a *de novo* interaction between CRBN and reporter protein.

To investigate if surface-edited CRBN is sufficient to induce neosubstrate degradation, we engineered a HiBiT tag sequence into the P1 site. Upon CRBN_HiBiT expression, we transfected a plasmid expressing LgBiT into the HAP1^*CRBN-*^cells. HiBiT tag structurally complements LgBiT, leading to an increase in luminescence. C-terminal fusion of HiBiT tag (C-term+DMSO) showed an enhanced luminescence compared to HiBiT insertion at the P1 site (P1+DMSO) across a range of LgBiT abundance (**Fig 1c**). Moreover, significant restoration of luminescence is observed when P1-HiBiT is treated with MLN4924 (P1+MLN4924) whilst minimal difference is observed between C-term+DMSO and C-term+MLN4924. Together, this data demonstrates that insertion of a HiBiT tag in P1 site of CRBN is sufficient to induce a neo-PPI which leads to the subsequent proteasomal-dependent degradation of the neosubstrate reporter, nanoLuc.

### High-throughput screening identifies gain-of-function insertions inducing GSPT1 degradation

To adapt the principles of neomorphic remodelling of an E3 ligase to a screening and discovery paradigm, we inserted a library of >125,000 nucleotide sequences into the P1 site of CRBN. We selected pomalidomide as a scaffold IMiD to activate CRBN and measured the cellular abundance of GSPT1 in the presence of each of the CRBN variants by pooled NGS-enabled screening (Marriott et al., 2002) (**Fig 2a**). We hypothesised that a suitable peptide insertion at the P1 site of CRBN could complement the interface between CRBN:pomalidomide and GSPT1, leading to a stable ternary complex formation and subsequent degradation of GSPT1 (Marriott et al., 2002; Matyskiela et al., 2016). GSPT1 protein abundance was measured by FACS to identify insertion sequences enriched in the GSPT1_low_ fraction. As GSPT1 plays an important role in eukaryotic translational termination (Chauvin et al., 2005), degradation of GSPT1 is observed to be cytotoxic (Matyskiela et al., 2016). Accordingly, a parallel phenotypic screen was performed to monitor the representation of individual CRBN mutant sequences for 10 days with continuous DMSO or pomalidomide treatment as a measure of cell fitness.

**Figure 2.**
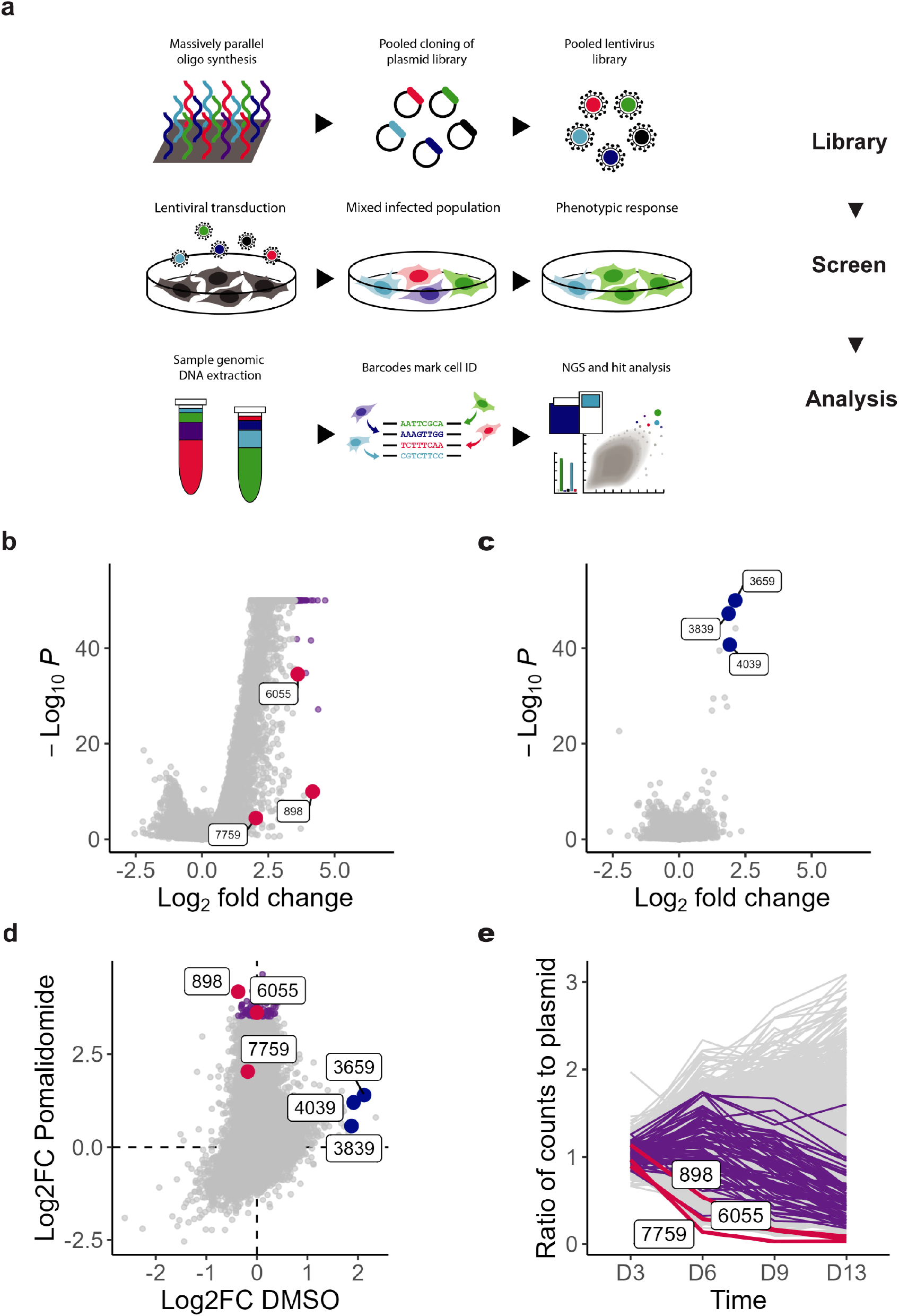
High-throughput GlueSEEKER screening identifies gain-of-function insertions required for GSPT1 degradation. a) Graphical representation of the process for high-throughput GlueSEEKER screening. A GlueSEEKER library (125K oligo pool) was computationally designed and cloned into lentiviral vector overexpressing CRBN at P1 site. HAP1^*CRBN-*^cells were then transduced with a pooled lentivirus library. Following selection and addition of spike-in controls, cells were treated with DMSO or pomalidomide. For GSPT1 degradation screening, FACS was used to isolate the top and bottom 15% GSPT1 populations following staining. A phenotypic outgrowth screen was performed in parallel with GSPT1 degradation screening, collecting cells immediately after selection (D3), and 3/6/10 days (D6/9/13) after DMSO/pomalidomide treatment. b) and c) Summary of pomalidomide-dependent (Fig 2b) and pomalidomide-independent (Fig 2c) GSPT1 degradation screening. Following NGS analysis, Log_2_ fold change (Log_2_FC) normalised counts in GSPT1 bottom 15% vs top 15% and −Log_10_P values were calculated. Of each AA sequence, the median of three independent barcodes is plotted (grey). A subset (purple) of the library is shown to have comparable performance with a known GSPT1 MGD CC-885 (black). Selected hits for arrayed validation are highlighted (red). d) Differential hit space between different treatments. Log_2_FC of median of three encodings of GSPT1 degradation screens are plotted as pomalidomide vs DMSO. More abundant hit space is observed with pomalidomide treatment. e) GSPT1 degrader hits show a time-dependent progressive reduction of counts. In phenotypic (cell fitness) screening performed in parallel to GSPT1 degradation screening, the hits identified in GSPT1 degradation screen are plotted as normalised to representation in library plasmids. GSPT1 hits (purple) show reduced representation compared to library (grey), consistent with CC-885 spike-in controls (black).

After collection, screening samples were analysed by deep sequencing and inserts on CRBN which induced a loss of GSPT1 determined by comparative analysis between gated populations. For GSPT1 degradation screening in the presence of pomalidomide (**Fig 2b**), a proportion of the library is observed to be enriched in the GSPT1_low_ fraction and a subset of the insertions show a comparable magnitude loss with a sample treated with a GSPT1-degrading molecule, CC-885, indicative of robust glue-like degradation. Without pomalidomide treatment (**Fig 2c**), both the portion of the library and the magnitude of enrichment in the GSPT1_low_ fraction are reduced, consistent with a previous report that IMiD-binding induces conformational rearrangements in CRBN which result in a shift in the functional output of the protein (Watson et al., 2022) (**Fig 2d**). Degrading variants of CRBN also display a corresponding kinetic loss of abundance in the cell fitness screen samples upon pomalidomide treatment (**Fig 2e**). Motif definition was robustly identified from the hits with a compelling similarity trend found (data not shown) and this was used to nominate the most bioactive sequences for further analysis. Taken together these results demonstrate that a high-throughput approach to effector protein remodelling can robustly identify novel sequence insertions that induce neomorphic function. The convergent properties of the active sequence space is indicative of a rationalizable mechanism that drives the interaction and is consistent with the aim of recruiting this same interface through small molecules.

### Surface remodelling of CRBN induces neosubstrate binding

To further understand the output of the screen, we first selected three sequence inserts from each screen arm (with and without pomalidomide treatment) to determine if GSPT1 degradation and the resulting anti-proliferative effect can be reproduced with alternative and individual measurements. Both pomalidomide-dependent and spontaneous GSPT1-degrading hits caused robust reduction of GSPT1 protein abundance compared to HEK293FT cells transfected with CRBN_WT (**Fig 3a**). Targeted, induced degradation of GSPT1 was also found to lead to a loss of cell fitness over time (**Fig 3b**), consistent with the expected effect of the loss of the protein to cells. Overall, there is a close correlation of the magnitude of loss in the individual assays and the effects measured by deep sequencing in the screening analysis, suggesting a robust and systematic readout.

**Figure 3.**
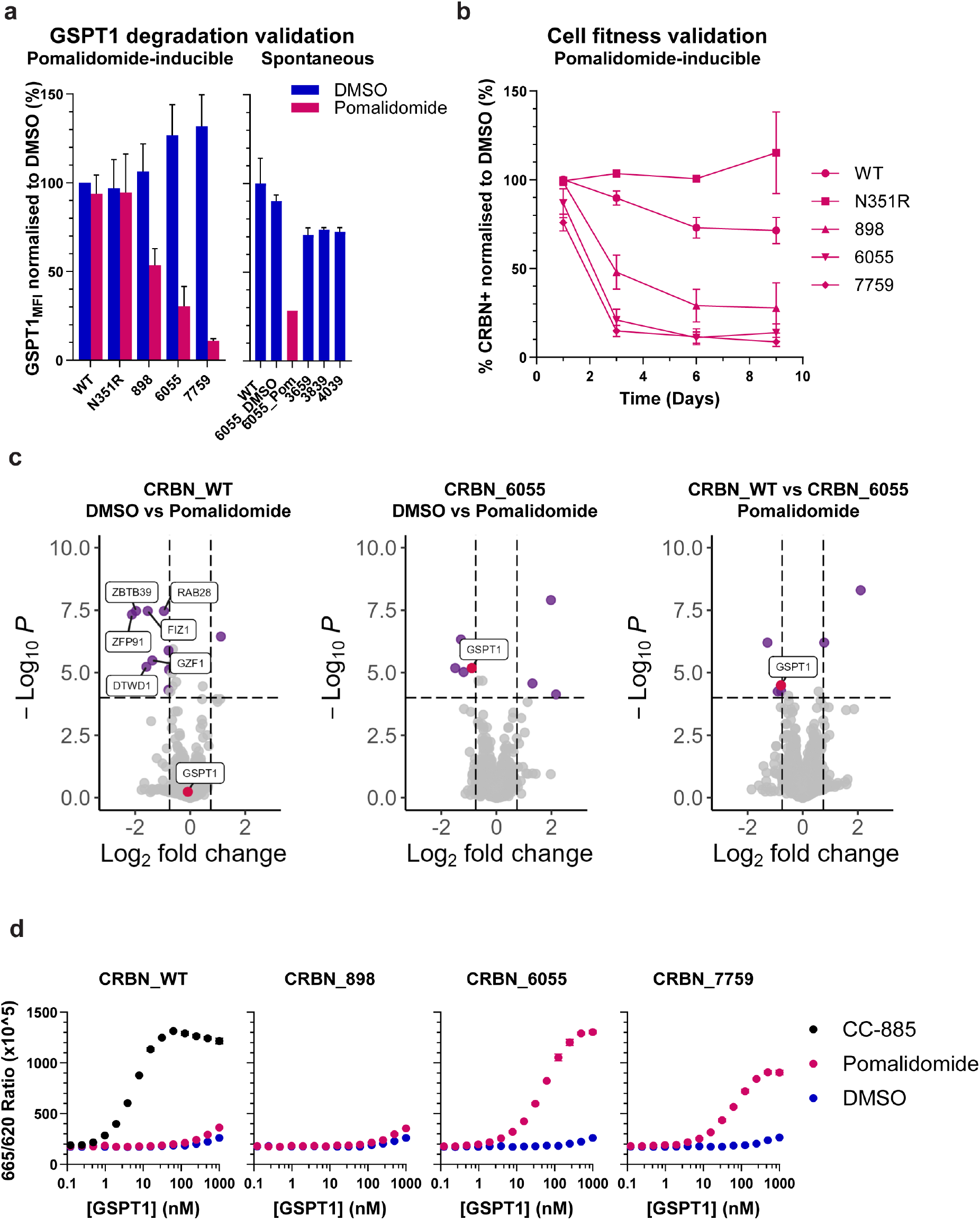
Validation of GSPT1 hits confirms precision editing of CRBN. a) Arrayed validation of GSPT1 degradation. Three pomalidomide-dependent (898/6055/7759) and three pomalidomide-independent (spontaneous hits; 3659/3839/4039) GSPT1 degrader hits are selected for further validation, together with CRBN_WT (WT), and a loss-of-function mutation (N351R (Van Nguyen et al., 2016)). For pomalidomide-dependent hits, HAP1^*CRBN-*^cells are transduced with lentivirus overexpressing CRBN and treated with DMSO or pomalidomide for 16 hours. Median fluorescence intensity (MFI) of GSPT1 is compared to CRBN_WT treated with DMSO. Significant reduction of GSPT1 is observed with GSPT1 hits treatment with pomalidomide, and the scale correlates with Log2FC from GSPT1 degradation screening. For pomalidomide-independent hits, HEK293FT cells were transfected with CRBN-overexpressing plasmids for four days. Median fluorescence intensity (MFI) of GSPT1 is normalised to CRBN_WT treated with DMSO. b) Validation of pomalidomide-induced reduction of proliferation of GSPT1 hits. During validation, upon DMSO or pomalidomide treatment, cells were harvested and the proliferation of CRBN-expressing cells monitored as the percentage of GFP+CRBN+ cells of total cell population over time. GSPT1 hits (898/6055/7759) show significantly reduced GFP+CRBN+ percentages when treated with pomalidomide, consistent with results from phenotypic screening. c) Hit 6055 degrades GSPT1 in pomalidomide-dependent fashion. TMT-global proteomics is used to identify neosubstrates targeted by hit 6055 upon 16-hour pomalidomide treatment. Compared with 6055 DMSO and WT POM, Hit 6055 shows significant reduction of GSPT1 abundance, whilst known neosubstrates of CRBN identified in POM WT vs DMSO WT but not GSPT1. d) Biochemical validation of ternary complex formation of GSPT1 hits. Recombinant CRBN (WT/898/6055/7759) were purified using a truncated backbone (Kroupova et al., 2024). Using an HTRF assay, ternary complex formation was detected between CRBN_WT, GSPT1 with CC-885 but not pomalidomide. With GSPT1 hits, robust ternary complex was detected with CRBN_6055 and CRBN_7759, consistent with observation from in cell validation (panels **a** and **b**).

We analysed whole cell lysates expressing the sequence insert 6055 and compared it to wild type CRBN in the presence or absence of pomalidomide using TMT mass spectrometry (**Fig 3c**). The most readily detectable proteins found to be depleted following the treatment with pomalidomide were overwhelmingly known or canonical substrates of this drug (left panel), indicating TMT-MS is sufficiently sensitive to detect neosubstrates targeted by CRBN. We found that GSPT1 is one of the most depleted proteins in cells expressing the CRBN variant 6055 (middle panel) and the protein was selectively depleted when compared with cells expressing wild type CRBN, which showed no loss of GSPT1 (right panel). Thus, the sequence insertion of 6055 is both necessary and sufficient to drive selective induced degradation of GSPT1 in the presence of the pomalidomide.

Finally, to confirm that GSPT1 degradation by surface-edited CRBN variants is a direct consequence of ternary complex formation between CRBN and GSPT1, we analysed the protein-protein interaction using recombinant proteins in an HTRF-based proximity assay (**Fig 3d**). When a CRBN protein (Kroupova et al., 2024) was purified as either wild type or one of three variants and co-incubated with recombinant GSPT1 we find that for the wild type proteins a complex formation is only induced by the molecule CC-885 but not pomalidomide or DMSO. In contrast, both CRBN_6055 and CRBN_7759 showed robust ternary complex formation with GSPT1 in the presence of pomalidomide, which was confirmed by analytical size exclusion chromatography (not shown) and with a response rate that was consistent with the degree of degradation observed in cellular assays for each insertion sequence.

### GlueSEEKER output enables novel MGD drug discovery

Following confirmation of direct GSPT1 targeting by surface-edited CRBN mutants, we next sought to identify novel GSPT1 MGDs using the chemical and geometric properties of the neomorphic CRBN. *In silico* structural modelling was used to establish a stable and novel interaction model of the modified CRBN and the GSPT1 neosubstrate. Whilst the overall architecture of the identified complex resembled that of the induced ternary association using extant molecular glues, several additional electrostatic interactions were predicted that rationalised the neo-PPI in the absence of small molecule binding (not shown). These new predicted hydrogen bonds and the spatial properties of the inserted loops were used to define new pharmacophores and conduct virtual screens that could identify molecules that recapitulate this ternary complex within the context of a WT CRBN:GSPT1 complex.

Overall, around 1500 compounds were selected for stocking, and tested for induction of a ternary complex using an homogenous time-resolved fluorescence (HTRF) based proximity assay. A number of these compounds were able to induce a stable complex between GSPT1 and CRBN (**Fig 4a**), which were confirmed in an orthogonal surface plasmon resonance assay (SPR; **Fig 4c**). The association was found to be robust, specific and pomalidomide-dependent for these early hit molecules, consistent with the results from pooled genetic screening (**Fig 4b** and counter screen data not shown). Weak binary interactions could be detected with CRBN, but not GSPT1, using ligand-observed NMR (**Fig 4d**).

**Figure 4.**
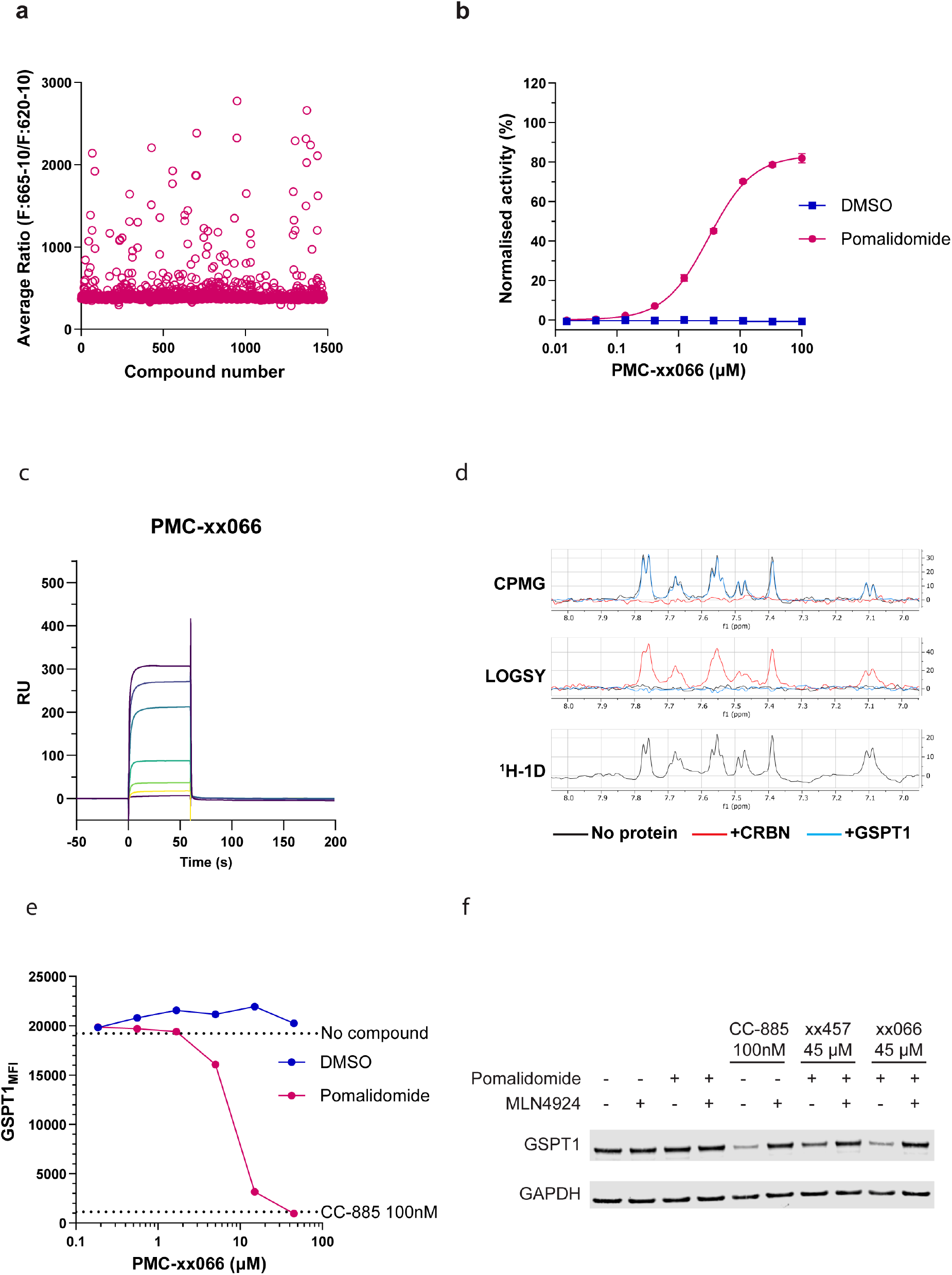
Small molecule discovery using GlueSEEKER. a) Screening results for ~1500 compounds identified by computational drug screening of around 5M structures *in silico* using an HTRF analysis of ternary complex formation for wild-type GSPT1 and CRBN protein. b) Dose response analysis for a leading hit from the screen in the presence or absence of pomalidomide. pomalidomide-induced ternary complex between CRBN and GSPT1 is observed with low micromolar activity. c) Biophysical analysis of ternary complex formation between purified recombinant CRBN and GSPT1 in the presence of fixed concentrations of pomalidomide, and leading hit PMC-XX066 by SPR. Rainbow traces show increasing concentrations of PMC-XX066. The estimated Kd under these conditions was 4 μM. d) Ligand observed NMR analysis of the low affinity binary interaction between leading hit PMC-XX066 and CRBN but not GSPT1. e) In-cell validation of GSPT1 degradation mediated by small molecule hit. HAP1^*CRBN-*^cells overexpressing CRBN_WT are treated with leading hit PMC-XX066 starting from 100 µM with or without pomalidomide. After 16h treatment, cells were harvested and median fluorescent intensity of GSPT1 staining in CRBN+ population plotted. pomalidomide- and PMC-XX066 dose-dependent reduction of GSPT1_MFI_is observed. f) Small molecule-mediated GSPT1 degradation can be rescued by on-pathway inhibition. HAP1^*CRBN-*^cells overexpressing CRBN_WT are treated with 1 µM pomalidomide, 100 nM CC-885, or 1 µM pomalidomide with leading hit PMC-XX066, in the presence or absence of 10 µM MLN4924. After 6h incubation, cells were harvested and whole cell lysates analysed by western blot. Significant reduction of GSPT1 observed for CC-885 and PMC-XX066 with pomalidomide and can be inhibited by MLN4924 treatment.

**Figure 5.**
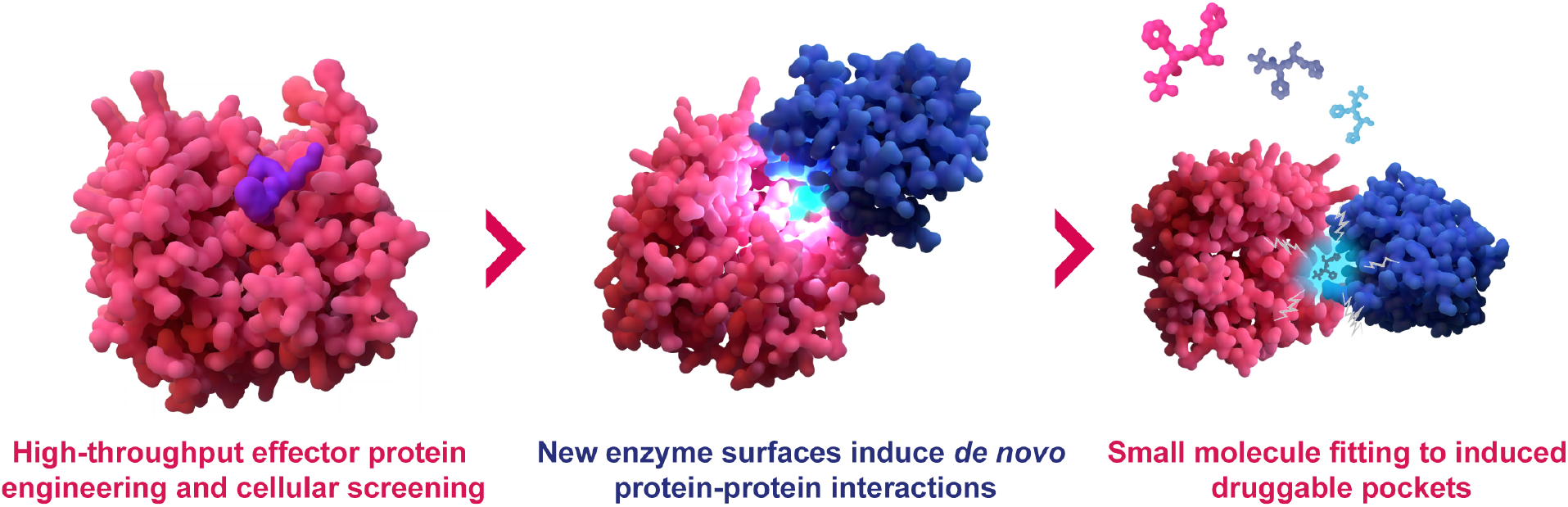
Graphical summary of the GlueSEEKER technology. High-throughput editing of effector protein surfaces enables gain-of-function screening to find new induced, *de novo* protein-protein interactions. Deconvolution, motif enrichment and computational modelling are used to recapitulate the interaction *in silico* and provide the blueprints to small molecule design and fitting to the induced pockets for molecular glue degrader discovery.

Promising hit molecules in the HTRF analysis were transitioned to a cellular assay for degradation. Cellular degradation of GSPT1 was observed for a substantial proportion of the *in vitro* active molecules (**Fig 4e** and not shown). Maximal GSPT1 degradation that approaches that of the benchmark molecule CC-885 was observed for several of the molecules, including PMC-XX066, albeit with higher DC_50_ concentrations. Degradation was robust and reversible with neddylation inhibitors (**Fig 4f**), indicating an on-mechanism induced degradation process is in operation in the cells.

Substantial similarity and clustering were also observed within the small molecule hit space, even despite the divergent library design after virtual screening, providing compelling hit matter for further development and an impressive demonstration of the end-to-end discovery of novel small molecules from a unique cellular protein engineering approach. Overall, these molecules operate as cellularly-active molecular glue degraders of GSPT1 in the presence of small molecule-treated CRBN. Computational docking of the molecules supports overlapping interaction moieties and spatial convergence with the hit inserted loops in the mutant variants of CRBN (not shown) and underlies the transformative approach to molecule screening or design by virtue of induced protein-protein interaction with this platform.

## DISCUSSION

PROTACs and MGDs are two major modalities of TPD showing enormous promise across the drug discovery landscape. Perhaps as a consequence of the functional similarities between the two modalities (namely, degradation), many of the screening tools applied so far to MGDs are based on heterobifunctional degrader discovery approaches - for example library diversification (Petzold et al., 2024), screening for cooperativity (Liu et al., 2023), or mass spectrometry (Steger et al., 2024). These tools possess good translation potential but crucially rely on extant ligand properties, have limited capacity for improved throughput and are generally less compatible with discovery of molecules beyond the small handful of demonstrated examples. Furthermore, it is now well understood that even the robust formation of a ternary complex is often not sufficient to induce degradation in cellular experiments (Baek et al., 2024). Instead, the first principles of protein-protein interaction and how this intersects with MGD chemical properties are needed to systematically discover these important new molecules. We have sought to address this gap by surveying the phenotypic consequences of effector protein editing at high-throughput in the physiological environment of cells and have shown neosubstrate activity of a key E3 ligase as a compelling demonstration for future and ongoing expansion. The prospect of widespread novel target discovery using degradation-enabled screening is also demonstrated by the cell fitness arms of the screening study which tightly converged with the GSPT1 target-centric approach.

Our experiments show that CRBN can be engineered at discrete sites to engage in neo-PPIs in cells and that with the right binding architecture, degradation is induced for neosubstrate and reporter proteins. Interestingly, we also found that targeted degradation of CRBN-bound reporter nanobodies was not observed, supporting the necessity to assess degradation-competent ternary complex formation in relevant physiological cell systems (Li et al., 2024). In the case of CRBN, one of the most studied and MGD-compatible E3 ligases, it is not clear if the G-loop degron sequence is either sufficient or even necessary to induce CRBN-mediated degradation (Petzold et al., 2024), and our approach presents the compelling opportunity to explore substrates beyond the canonical and predicted subset to further define this. The observation in this study that mutation-induced neomorphic activity of CRBN was potentiated by co-treatment with an IMiD supports the notion that CRBN requires conformational activation, but clearly this is not the only mode of activity given its endogenous role (Ichikawa et al., 2024). In this context, the observation that a small but notable hit space in our screens was found not to require pomalidomide to induce GSPT1 degradation is highly intriguing and will be important to further characterise to understand the potential for non-IMiD molecular glue discovery with CRBN. Furthermore, this observation supports the strong expectation that the molecules reported in this study can be refined and optimised to dispense with the IMiD co-treatment, offering the appealing prospect of a new non-IMiD ligand class to engage and degrade both GSPT1 and other important neosubstrate proteins via CRBN, potentially circumventing the known liabilities of this class of drug.

A recent study examining mutations of the E3 ligase, KBTBD4, found that neosubstrate degradation of the CoREST complex was triggered by medulloblastoma-associated insertions in the Kelch pocket of the ligase (Chen et al., 2022). Moreover, the electrostatic interactions that conferred stability of the complex were exquisitely mimicked by the small molecule UM171, redefined as a *bona fide* MGD (Chen et al., 2022; Xie et al., 2024; Yeo et al., 2025). Thus, induction of protein-protein interactions by gain-of-function insertions can be both structurally and functionally mimicked by small molecules. We have expanded on this principle in the current study using a forward genetic screening approach to induce novel and biologically productive PPIs which can be used to rationally inform small molecule drug discovery. We have successfully extended this approach to other E3 ligases (not shown), confirming the approach is generalisable beyond the CRBN example reported on here. By further targeting functional hotspots on E3 ligases to induce neo-PPI, our GlueSEEKER technology enables a high-resolution understanding of the physical and chemical requirements of neo-PPIs suitable for novel MGD therapeutics development and provides high-fidelity training material for machine learning of generative PPIs and chemical design.

## METHODS

### Functional evaluation of re-constituted CRBN constructs

CRBN overexpression plasmids encoding CRBN_WT, CRBN_P1-HA, CRBN_P2-HA and CRBN_P1-HiBiT were cloned into second generation lentivirus packaging plasmids by InFusion reaction following manufacturer’s instructions. CRBN expression was reconstituted in HAP1^*CRBN-*^cells (Horizon Discovery) following lentivirus infection at MOI=0.3 and selected by 1 µg/mL puromycin. For dBET6-mediated degradation assay, cells were seeded in 24-well plates one day before compound treatment followed by incubating with 0.5% DMSO, 100 nM dBET6 or 100 nM dBET6 + 1 µM MLN4924 for 4 hours and 18 hours. After incubation, cells were lysed with RIPA buffer containing 1x protease inhibitors cocktail (Roche). Protein concentration was measured by BCA assay and 10 µg whole cell lysates were analysed by 3-8% NuPAGE™ Tris-Acetate (for BRD4) and 4-12% NuPAGE™ Bis-Tris gels (for CRBN and GAPDH) and transferred by 0.2 µm nitrocellulose membrane (Bio-Rad). Membranes are developed by Odyssey M Imager (LICORbio).

### Co-immunoprecipitation

CRBN expression was reconstituted in HAP1^*CRBN-*^cells as described in compound treatment. After selection by puromycin, cells were superinfected with recombinant lentivirus encoding EGFP-HA nanobody and blasticidin resistant gene (Zhao et al., 2019). Following blasticidin selection (10 µg/mL), 4×10^6^ cells were seeded overnight and harvested by trypsin. Anti-GFP immunoprecipitation was performed using GFPTrap magnetic agarose following manufacturer’s instruction (Chromotek). After overnight rotating incubation, magnetic agarose were washed and eluted with 5x equivalent concentration. 1% input and 5% IP fractions are analysed by western blot.

### Luciferase assay

CRBN expression is reconstituted in HAP1^*CRBN-*^cells as previously described. Following puromycin selection, cells were seeded in 96-well plates and transfected with plasmid encoding LgBiT using Lipofectamine2000. 48 hours post transfection, cells were treated with 0.5% DMSO or 10 µM MLN4924 for 6 hours before luminescence detected using Nano-Glo® HiBiT Lytic reagent (Promega).

### GlueSEEKER Screening

Effector proteins (e.g. CRBN) insertion sites are nominated by modelling and *in silico* energetic predictions and insertion libraries designed and encoded for synthesis by oligo printing (Twist Bioscience). Libraries of lentivirus are produced and cells are infected and selected at over 300X representation ahead of screening initiation, sampling and treatments. Sample collection is by high-throughput FACS and genomic material is extracted similarly to previous reports (Cross et al., 2016). Data processing and analysis: DEseq2 (v1.42.1) was used to normalise counts between samples and detect differential representations of sequences between the GSPT1 sorted bins (Love et al., 2014). The resulting significance values were adjusted to control for FDR using the Benjamani-Hochberg procedure. Amino acid sequences which were identified as significantly enriched in the GSPT1^Low^ across multiple DNA encodings were called as potential GSPT1 degrading sequences. For the dropout screen, NGS counts were normalised with DEseq2, and we calculated the mean ratio of counts at each time point to the plasmid for each designed insert sequence. Graphs were plotted with EnhancedVolcano (v1.20.0) or ggplot2 (v3.5.1) (Blighe, 2018; Wickham, 2009). For arrayed validation, individual sequences were re-cloned and introduced into cells either by lentivirus or by transient transfection. Selected variants were analysed for protein abundance or cell fitness by flow cytometry, western blotting or metabolic cell health assays.

*Ternary complex formation HTRF assay with CRBN mutants* Complex formation between CRBNΔHBD WT and mutants with GSPT1_300-469_C-Avi-biotin in the presence of DMSO or fixed concentrations of pomalidomide or CC-885 was assessed using HTRF. Compounds or DMSO were dispensed into a 384 well plate using a Tecan D300e. CRBN constructs were dispensed into the 384-well plate. GSPT1_300-469_C-Avi-biotin was titrated with Streptavidin-d2 (Revvity) and added to the 384 well plate along with Anti-MBP-Tb (Revvity). The plate was mixed and incubated for 180 minutes. 665nm/620nm ratio were measured using a CLARIOStar plate reader (BMG).

### Analytical Size Exclusion Chromatography

Size exclusion chromatography was carried out using a Superdex 200 Increase 10/300 column in 50 mM HEPES pH 7.5, 250 mM NaCl, 2% glycerol and 1 mM TCEP using an ÄKTA Avant (Cytiva) at a flow rate of 0.5 mL/min. Equimolar concentrations of CRBNΔHBD-6055 and pomalidomide, CRBNΔHBD-6055, GSPT1_300-469_C-Avi and pomalidomide, or GSPT1_300-469_C-Avi alone were analysed. Eluted fractions were analysed by SDS-PAGE.

### Sample dissolution, TMT labelling and Reverse-Phase fractionation

The cell pellets were resuspended in lysis buffer containing 100 mM triethylammonium bicarbonate (TEAB, Sigma), 10% isopropanol, 50 mM NaCl, 1% SDC with nuclease and protease/phosphatase inhibitors, followed by 15 min incubation at RT. Later protein concentration was estimated using Bradford assay according to manufacturer’s instructions (BIO-RAD-Quick start). 90 ug of total protein were reduced and alkylated with 2 ul of 50 mM tris-2-caraboxymethyl phosphine (TCEP, Sigma) and 1ul of 200 mM iodoacetamide (IAA, Sigma) respectively for 1 hour. Then protein samples were digested overnight at 37°C using trypsin solution at ratio protein/trypsin ~ 1:30. The next day, protein digest was labelled with the TMTpro reagents (Thermo Scientific) for 1 hour. Later the reaction was quenched with 4 μL of 5% hydroxylamine (Thermo Scientific) for 15 min at room temperature (RT). All the samples were mixed and acidified using formic acid for SDC removal. Later the mix was dried with speed vac concentrator. The dry TMT mix was fractionated on a Dionex Ultimate 3000 system at high pH using the X-Bridge C18 column (3.5 μm, 2.1×150 mm, Waters) with 90 minute linear gradient from 5% to 95% acetonitrile containing 20 mM ammonium hydroxide at a flow rate of 0.2 ml/min. Peptides fractions were collected between 20-55 minutes and were dried with speed vac concentrator. Proteome fractions were reconstituted in 0.1% formic acid for liquid chromatography tandem mass spectrometry (LC–MS/MS) analysis.

### LC-MS/MS

Peptide fractions were analysed on a Vanquish Neo UHPLC system coupled with the Orbitrap Ascend (Thermo Scientific) mass spectrometer. Peptides were trapped on a 100 μm ID X 2 cm microcapillary C18 column (5 µm, 100A) followed by 90 minutes elution using 75 μm ID X 25 cm C18 RP column (3 µm, 100A) at 300 nl/min flow rate. A Real Time Search (RTS)-MS3 method was used for the analysis; The MS1 spectra were acquired in the Orbitrap (R=120K; scan range 400-1600 m/z; AGC target = 400000; maximum IT = 251 ms) and the MS2 spectra in the Ion Trap (isolation window 0.7 Th; collision energy of 30%; maximum IT = 35 ms; centroid data). For RTS, Trypsin/P digestion was selected using static cysteine carbamidomethylation and TMTpro modification on lysine and peptide N-terminus. The search was conducted for a maximum of 35 ms with the following thresholds: Xcorr =1.4, dCn = 0.1, precursor ppm 10, charge state = 2. MS3 spectra were collected in the Orbitrap (R= 45K; scan range 100−500 m/z; normalized AGC target = 200%; maximum IT = 200 ms; centroid data. Phospho fractions were subjected to MS2 analysis without RTS. Data processing: The Proteome Discoverer 3.0. (Thermo Scientific) was used for the processing of CID tandem mass spectra. The SequestHT search engine was used, and all the spectra searched against the Uniprot Homo sapiens FASTA database (taxon ID 9606 - Version June 2024). All searches were performed using a static modification of TMTpro (+304.207 Da) at any N-terminus and lysines and methylthio at cysteines (+45.988 Da). Methionine oxidation (+15.9949 Da), phospho (+79.966 Da on serine, threonine and tyrosine) and deamidation (+0.984) on ssparagine were included as dynamic modifications. Mass spectra were searched using precursor ion tolerance 20 ppm and fragment ion tolerance 0.5 Da. For peptide confidence, 1% FDR was applied, and peptides uniquely matched to a protein were used for quantification (Langmead and Salzberg, 2012).

### Mass spectrometry

Expression data was imported with MSnBase (v2.28.1), filtered to only include Master Proteins and global median normalised |(Gatto and Lilley, 2012). Statistical testing was performed by the Limma R package (v 3.58.1) to fit a linear model per protein, which were moderated by empirical Bayes smoothing (Ritchie et al., 2015). The resulting significance values were adjusted to control for FDR using the Benjamani-Hochberg procedure. Graphs were plotted with EnhancedVolcano (v1.20.0) (Blighe, 2018).

### Ternary complex formation assay by HTRF and compound screening

The compound library was initially screened at 100 μM and 30 μM for their ability to promote ternary complex between MBP-CRBNΔHBD and GSPT1_300-469_C-Avi-biotin in the presence of 5 μM pomalidomide. Briefly, compounds were stamped out into 384-well plates. MBP-CRBNΔHBD/GSPT1_300-469_C-Avi-biotin/pomalidomide were dispensed onto the compounds, mixed and incubated for 30 minutes. Anti-MBP-d2 donor and Streptavidin-Tb (Revvity), were dispensed in to the plate, incubated for 60 minutes. 665nm/620nm ratio was measured using a CLARIOStar plate reader (BMG). For compounds dose responses, compounds were dispensed into a 348-well plate using a Tecan D300e. The assay was then performed as for the dual point screen.

### Surface Plasmon Resonance (SPR)

SPR experiments were carried out on a Biacore S200 (Cytiva) operating at 25°C, flow rate 30 μL/min, in 1X HBS-P+ (Cytiva), 0.5 mM TCEP, 2% DSMO. GSPT1^300-469^C-Avi-biotin was immobilised onto the surface of an SA Chip (Cytiva) to ~250-500 resonance units (RU). Serial dilutions (50 μM-70 nM) of CRBNΔHBD (Bailey et.al 2024) were injected in the presence of 100 μM pomalidomide and 100 μM compound. Kds were determined from steady-state analysis using a Langmuir 1:1 binding model. Exclusion of either pomalidomide or MGs results in loss of binding response (data not shown).

### Nuclear Magnetic Resonance (NMR) Spectroscopy

NMR experiments were carried out on a Bruker Avance DRX500 (Ultrashield) spectrometer equipped with 5 mm HCN room temperature probe at 5°C in PBS pH 7.5, 5% d6-DMSO. Samples contained 500 μM of MG or pomalidomide in the presence or absence of 10 μM CRBN-ΔHBD or GSPT1_300-469_. Binary interactions between compounds and target proteins were assessed using either Water-Ligand Observed via Gradient SpectroscopY (WaterLOGSY) or Carr-Purcell-Meiboom-Gill (CPMG) based methods.

## ACKNOWLEDGEMENTS

The authors thank all the team members at PhoreMost for helpful comments and contributions during the preparation of this manuscript and the flow cytometry facility of the Babraham Institute and CRUK Cambridge for support with FACS and mass spectrometry, respectively. We thank the Biophysics and NMR Facilities in the Department of Biochemistry, University of Cambridge for access to instrumentation.

